# RegulomeXplorer: Interactive exploration of drug effects on subcellularly resolved proteomes

**DOI:** 10.64898/2026.06.29.735319

**Authors:** Matthias Uiberacker, Thomas Iellici, Elena Afanaseva, Samuel M. Meier-Menches, Jürgen Zanghellini

## Abstract

Mass spectrometry-based proteomics allows the quantification of drug-induced changes in protein abundance. However, the integration of perturbation data across subcellular compartments remains a challenging bottleneck. Here, we present `RegulomeXplorer`, a web-based tool for automated processing and interactive exploration of subcellular compartment-resolved proteomics data. `RegulomeXplorer` employs MaxQuant output files to determine differential protein regulations upon drug perturbation, performs functional enrichment analysis, and visualizes enriched terms on a two-dimensional cytoplasmic-nuclear plane, called regulome. The data visualization by means of regulomes allows to simultaneously assess the magnitude of drug perturbation effects within separate subcellular compartments as well as the contribution of regulated proteins to the position of each enriched term in the regulome plane. We validated `RegulomeXplorer` against previously published, manually curated regulome analyses. It was then applied on subcellular compartment resolved breast cancer cell line proteomes, revealing drug- and cell-line-specific responses to Doxorubicin and Taxol, both in line with their described mode of action. `RegulomeXplorer` provides an accessible workflow for interpreting compartment-resolved perturbation proteomics and generating mode of action hypotheses in drug-response studies. `RegulomeXplorer` is freely available without registration at https://chemnettools.anc.univie.ac.at/RegulomeExplorer/.

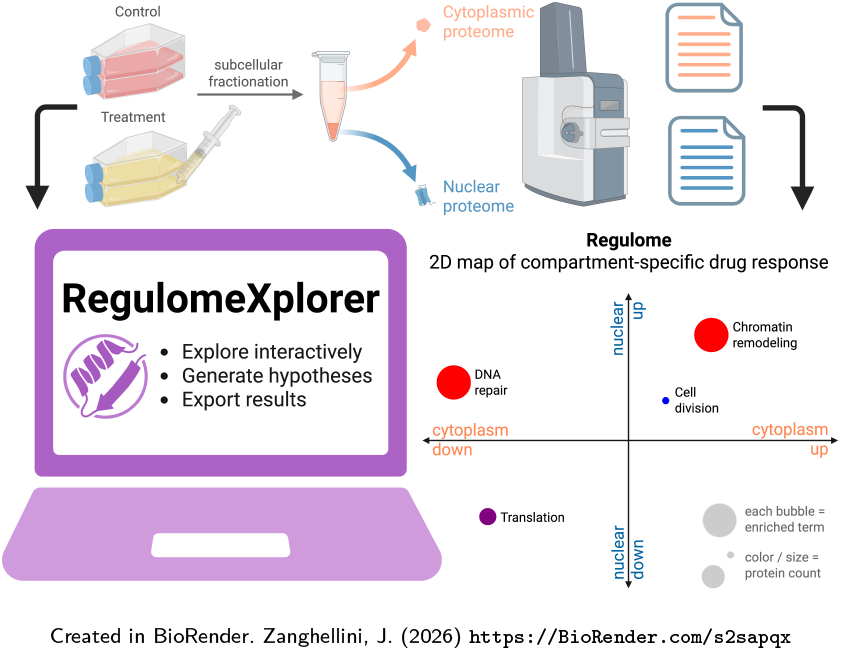

## Introduction

Mass spectrometry-based proteomics has become a central technology for drug discovery [1, 2], enabling the identification of drug targets [3, 4], the characterization of cellular response signatures [5], and the discovery of biomarkers associated with drug response [6]. In perturbation proteomics, pharmacological treatment shifts a cellular system away from homeostasis, and the resulting changes in protein abundance provide a molecular signature that can reveal the mode of action (MoA) of a compound. This signature-driven approach relies on robust experimental and statistical workflows to control technical and biological variation and ensure high-quality perturbation profiles [7]. Automated sample preparation and multiplexing now allow such perturbation profiles to be generated at large scale, including datasets comprising millions of drug-protein associations [8] across compounds, cell lines [5], and dose ranges [9]. However, transforming differential protein abundance data into coherent biological insight and testable MoA hypotheses remains a major analytical and computational bottleneck, particularly for experimental proteomics laboratories without dedicated computational support.

Cellular adaptation to pharmacological treatment unfolds over several hours to days and involves coordinated changes in signaling pathways, gene regulation, protein abundance, and subcellular localization [6]. Functional enrichment analysis is widely used to convert regulated protein lists into interpretable biological categories, for example by identifying gene ontology (GO) terms or pathways that are overrepresented among regulated proteins [10, 11]. Web resources such as DAVID [12, 13], STRING [14], and KEGG-based pathway analyses [15] support this step and are widely used in proteomics. However, when cytoplasmic and nuclear fractions are analyzed as separate protein lists, the relationship between the two compartment-specific responses is lost; when they are combined into a single list, compartment-specific regulation is obscured.

A related limitation arises from the experimental design of many perturbation proteomics studies, which often rely on whole-cell lysates because they are experimentally straightforward and compatible with automated sample preparation workflows [9, 8, 5]. Whole-cell measurements collapse compartment-specific regulation into a single abundance profile, although proteins often exert their biological functions in defined subcellular locations. Nucleocytoplasmic fractionation separates two major functional spaces of eukaryotic cells, the cytoplasm and the nucleus, and has been used to study compartment-resolved proteomes [16] and drug-induced perturbations [17, 18]. When combined with functional enrichment analysis, such data can reveal whether biological processes are regulated globally, selectively in one compartment, or in opposite directions between cytoplasm and nucleus. However, extracting these relationships from differential proteomics data currently requires multiple disconnected analysis steps, manual data handling, and custom visualization.

To address this gap, we developed `RegulomeXplorer`, an integrated web-based data processing and visualization workflow for nucleocytoplasmic label-free quantification (LFQ) perturbation proteomics. `RegulomeXplorer` converts MaxQuant result files into compartment-specific differential-expression results, performs functional enrichment analysis, and generates interactive two-dimensional perturbation plots, termed regulomes. In these plots, each point represents an enriched GO term or pathway, while its position summarizes the regulation of the associated proteins in the cytoplasmic and nuclear fractions. This representation allows users to distinguish biological processes that are co-regulated in both compartments from those showing compartment-specific or opposing responses, thereby supporting hypothesis generation about drug MoA, cellular stress responses, and potential resistance-associated processes. Both the web application and source code are publicly available under an MIT license at https://chemnettools.anc.univie.ac.at/RegulomeExplorer/, and https://github.com/chemnet-univie/regulome-xplorer, with documentation and example datasets to facilitate adoption. We demonstrate the utility of this framework by applying it to perturbation datasets comprising established anticancer agents and investigational compounds across multiple cancer cell lines.

## Methods

### Experimental datasets

`RegulomeXplorer` was evaluated using previously published nucleocytoplasmic perturbation proteomics datasets.

The first dataset comprised SW480 colon cancer cells treated for 24 h with the investigational gold compound [Au(CCON)Cl_2_] (CCON = 2-benzoylpyridine; Au003/RBA29) or vehicle control, with six biological replicates per condition [19].

The second dataset comprised four breast cancer cell lines representing major clinical subtypes: MCF-7 (Luminal A), BT-474 (Luminal B/HER2+), MDA-MB-231 (triple negative), and SK-BR3 (HER2+) [20]. Cells were treated for 72 h with cell-line-specific IC_50_ concentrations of Doxorubicin (Dox) or Taxol (Tax), or with vehicle control. Three biological replicates were available per condition.

For both datasets, cytoplasmic and nuclear fractions had been generated by nucleocytoplasmic fractionation and analyzed by label-free LC–MS/MS as described in the original studies [19, 20]. Raw data had been processed with MaxQuant [21] against the UniProt Swiss-Prot human database [22], using a peptide and protein false discovery rate of 0.01 and match-between-runs enabled. The resulting MaxQuant protein-group tables containing UniProt identifiers and LFQ intensity values were used as input for RegulomeXplorer.

`RegulomeXplorer`’s computational framework

### Architecture and implementation

`RegulomeXplorer` is implemented in Python *3.11*, uses rpy2 R-bridge with R *4.3* for expression analysis and Dash *4.0.0* and Plotly *6.6.0* for UI and visualization. Data are stored in a MySQL database (version *8.4.5*) accessed via SQLAlchemy *2.0.40*. The application is deployed on a standalone server running on Ubuntu Linux *24.04.4 LTS*. The source code is released under the MIT license at https://github.com/chemnet-univie/regulome-xplorer.

A Conda environment file specifying all dependencies and exact versions is provided. The overall workflow from data import to regulome visualization is summarized in Figure 1.

**Fig. 1:**
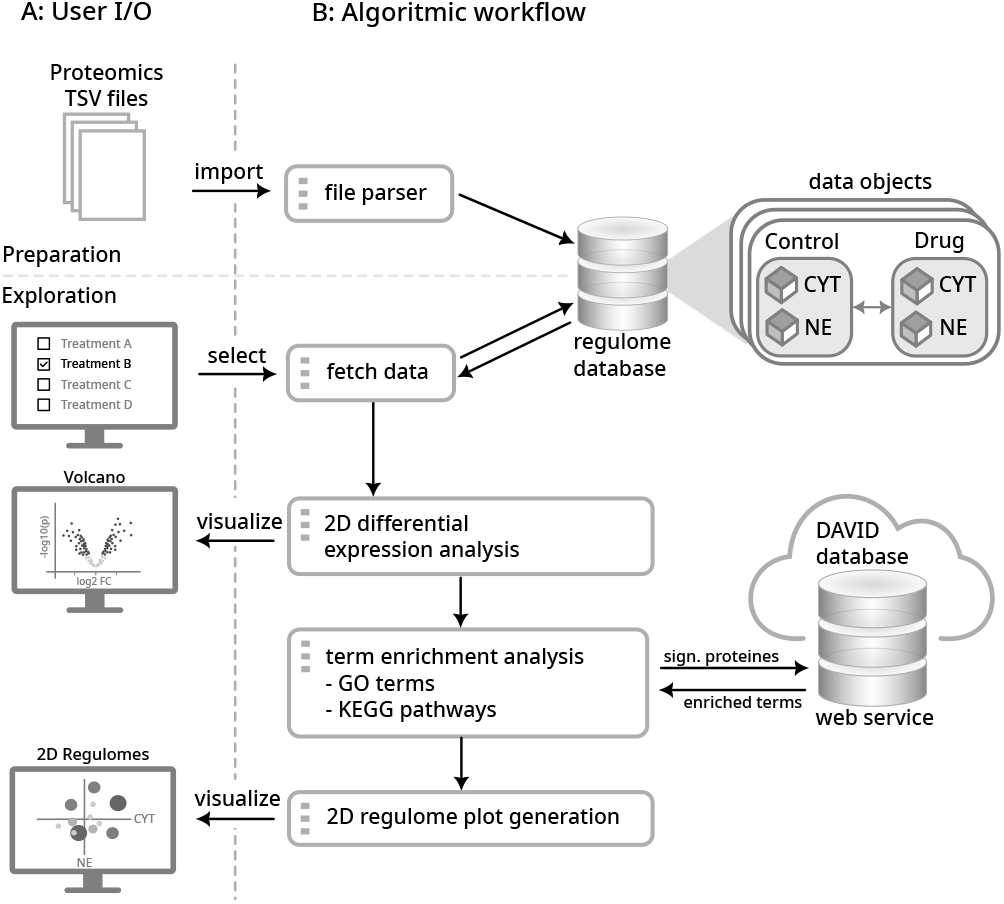
Proteomics workflow: (A) User interaction: Max Quant TSV output files can be imported via python script. (B) Algorithmic workflow: User selects data to explore. In turn data tables, 2 volcano plots for both cell fractions and 4 regulome plots for biological processes, molecular functions and cellular compartments (GO terms) as well as KEGG pathways are generated and interactively visualized.

### Data import and preprocessing

Input datasets are provided as tab-separated MaxQuant LFQ output files containing UniProt Accession numbers and replicate-specific intensity values for cytoplasmic and nuclear fractions. Upon upload, files are parsed, validated for structural consistency and converted into a standardized internal data representation. At minimum, protein identifiers and LFQ intensity columns for control and treatment replicates per fraction are required; see detailed specifications in the Supplementary Information, section *Input File Format*. Data are serialized as structured JSON objects to support efficient querying and interactive visualization, and stored in the relational database together with associated metadata.

Missing intensity values are retained as NA. No imputation is performed - the downstream statistical analysis operates on available replicate measurements only. The default minimum required number of measured values is set to 5 for one condition (treatment or control) and 3 for the other to achieve a reasonable tradeoff between statistical power and usable proteome size. The thresholds can be adapted depending on the number of measured replicates in a given dataset. Proteins with fewer values will be discarded for differential expression analysis.

### Dataset selection

Datasets can be selected for analysis via the web interface. Upon selection, fraction-specific treatment and control data are loaded from the database into a structured internal representation separating cytoplasmic and nuclear measurements for downstream statistical analysis (Fig. 1).

### Dataset visualization

The proteome data is visualized in a data table, where each row corresponds to a measured protein. The average log2 LFQ intensity values are given for treatment and control for both fractions. The table is sortable and can be filtered for e.g. certain protein or gene ids (Fig. 2 A).

**Fig. 2:**
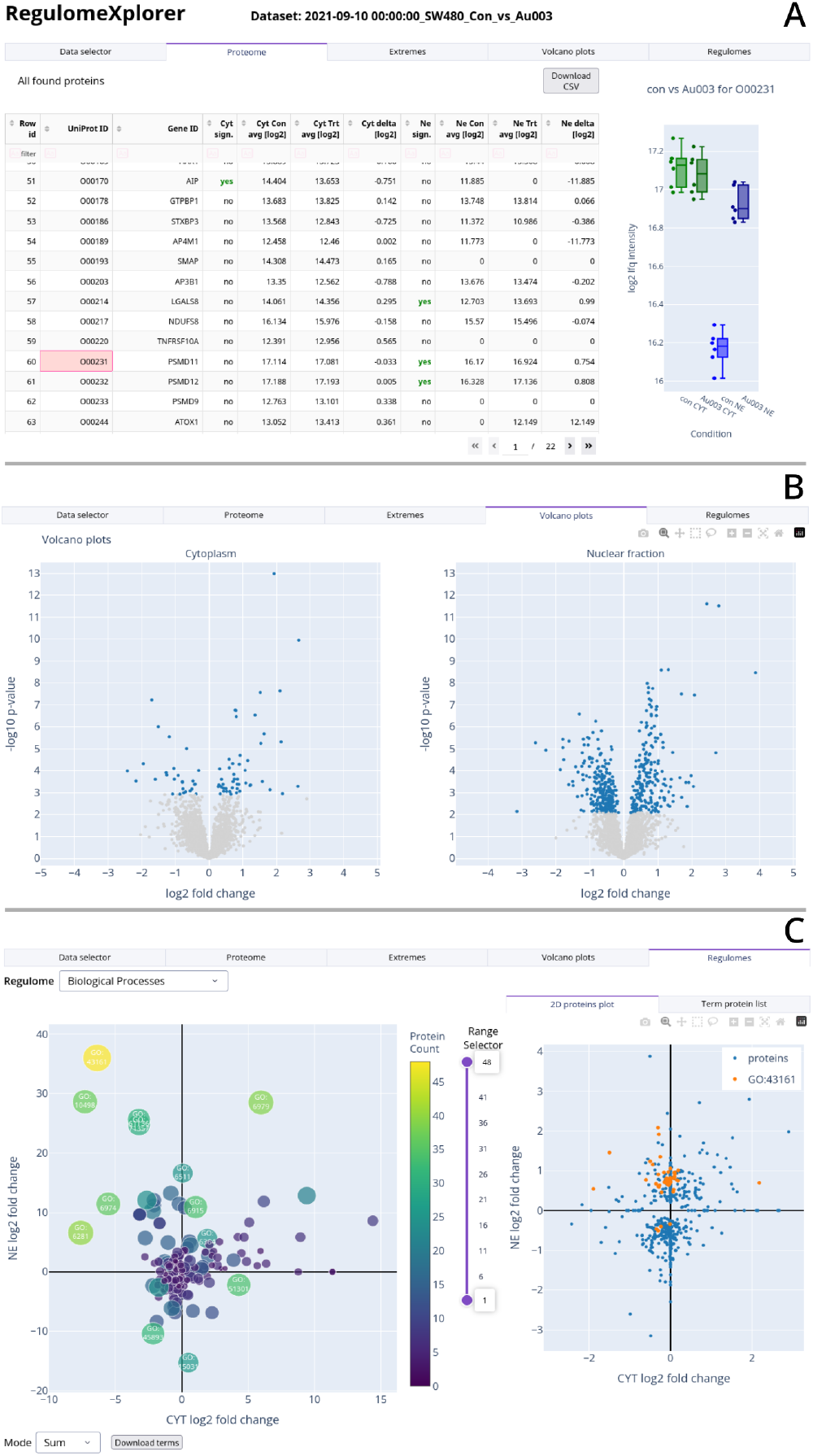
Main sections of the user interface: (A) Filter- and searchable proteome table. Intensity values are given for control and treatment for both fractions. Zeros are shown for missing values. Upon row selection, all individual LFQ intensity values are shown in detail in the adjacent plot. (B) Volcano plots (left: cytoplasmic fraction, right: nuclear fraction). (C) Regulome plot of category *biological processes*: Each term is represented as a circle on a two-dimensional (2D) plane. In *Sum* mode (as shown here) CYT and NE axes represent the summed fold-change for the cytoplasmic and nuclear fraction, respectively. In *Average* mode the circle is positioned at the average instead. Color and size represent the number of proteins per term. Upon selecting a term, the corresponding proteins are highlighted in the adjacent 2D expression plot.

### Differential expression analysis

For both the cytoplasmic and nuclear fraction an individual differential expression analysis is performed using the **limma** (Linear Models for Microarray Data) R-package from Bioconductor via the Python-R bridge. The false discovery rate (FDR) is set to 5%. Volcano plots are generated side by side for both fractions showing the significantly expressed proteins (Fig. 2 B).

### Gene ontology term enrichment analysis

The sets of statistically significant protein regulations of both subcellular fractions are combined to one list of unique proteins. This list serves as the input for a GO term enrichment analysis. To access curated and regularly updated protein and GO annotation data, `RegulomeXplorer` interfaces with the publicly available DAVID Bioinformatics web services (National Institutes of Health). Our algorithm connects via API to the DAVID web-service, inputs the list of significantly regulated protein IDs (uniprot accession number) and queries for an enrichment analysis of GO terms and KEGG pathways. DAVID performs the term enrichment analysis using a one-tail Fisher Exact p-value (named EASE score - see https://davidbioinformatics.nih.gov/documentation for documentation). We use an EASE score threshold of 0.05. The minimum number of significant genes per term is set to 2. The API response contains the enriched GO terms for biological processes (BP), molecular functions (MF) and cellular compartments (CC) as well as enriched terms of KEGG pathways.

### Data visualization as 2D regulome plots

The graphical representation of regulated terms within the 2D space of cytoplasm and nuclear fraction has been referred to as regulome plot [19]. Each term will be represented as a data point on a 2D scatter plot. The sum (or average) of fold-changes per fraction form the x and y coordinates for a term, where x stands for fold-change in the cytoplasmic fraction (CYT) and y for fold-change in the nuclear fraction (NE). The number of significantly regulated proteins of a term is represented by the size of the data point and its color (Fig. 2 C). Four individual regulome plots will be prepared for the four categories: BP, MF, CC and KEGG.

The two subcellular axes of the regulomes encompass four quadrants, which are of biological importance. The top-right quadrant represents terms or pathways, whose proteins are upregulated in both cytoplasmic and nuclear fractions, while the bottom-left quadrant represents terms or pathways that are both down-regulated. The other two quadrants show terms that are upregulated in one and downregulated in the other fraction and therefore indicate terms or pathways that undergo nucleocytoplasmic shifts or translocations.

Terms of regulomes can optionally be visualized according to summed or averaged fold-changes of their corresponding proteins. While averaging normalizes the fold-changes of protein groups in term objects, summed fold-changes do not. The summed approach produces stronger contrasts based on the assumption that proteins corresponding to a particular biological process may act in concert but not necessarily involve large average changes in abundance. See Supplementary Information, section *Regulome visualization mode* and Supplementary Fig. S3 for details.

The plot can be further customized by selecting the term size range to e.g. exclude small terms or terms with a large number of proteins.

When clicking on a term, the corresponding protein distribution of the selected term is highlighted in the adjacent 2D plot showing all significant proteins. The list of the proteins can be viewed and exported for the selected term.

## Results

To evaluate `RegulomeXplorer`, we first assessed its agreement with a manually curated regulome analysis [19] and then used it to analyze breast cancer perturbation datasets for compartment-resolved comparison of drug responses across cell lines and compounds [20].

### RegulomeXplorer reproduces manually curated regulomes

Relative to the manually curated reference analysis, `RegulomeXplorer` generated cytoplasmic and nuclear volcano plots with identical data points and only minor differences in significance classification (Supplementary Fig. S1). The corresponding GO BP and GO CC regulomes showed largely conserved term composition and positioning in the cytoplasmic–nuclear response space (see 8.2 and Supplementary Fig. S2).

The minor differences in significance classification (Supplementary Fig. S1) and regulome term placement (Supplementary Fig. S2) were expected from the comparison design. The published workflow used left-censored imputation followed by differential expression analysis with permutation-based significance analysis of microarrays (SAM) in Perseus, whereas the default `RegulomeXplorer` workflow avoids imputation and uses linear models for microarray data (LIMMA). To make the comparison as close as possible, we used the original imputed input table and applied SAM from the `siggenes` Bioconductor package.

Remaining differences in significance classification can therefore be attributed mainly to implementation-specific differences between the Perseus and `siggenes` versions of SAM, including the random permutation procedure and false discovery rate (FDR) threshold estimation. Because regulomes are generated from proteins classified as significantly regulated, small upstream differences can propagate into the enrichment analysis, slightly altering term size and position. DAVID knowledge-base updates may further affect GO annotations and term membership.

Together, these results demonstrate that `RegulomeXplorer` automates the generation of compartment-resolved regulomes.

### RegulomeXplorer reveals drug- and cell-line-specific responses

We next applied `RegulomeXplorer` to a published perturbation dataset [20] comprising cytoplasmic and nuclear proteomes from four breast cancer cell lines: MCF-7 (Luminal A), BT-474 (Luminal B), SK-BR3 (HER2+), and MDA-MB-231 (triple-negative), representing major clinical tumor subtypes [23]. Cells were treated with half-maximal growth inhibitory concentrations of Dox or Tax, two chemotherapeutic agents with distinct mechanisms of action targeting DNA topology and microtubule dynamics, respectively [24, 25, 26].

All datasets consist of 3 biological replicates with 2 technical replicates each. Therefore, we adapted the acceptance criterion for the analysis to have measured values for at least 2 biological samples with their 2 technical replicates in both conditions. The positions of biological and technical replicates were passed to LIMMA to correctly differentiate between them statistically.

Across all four cell lines, the Dox regulomes showed perturbation patterns in line with the known MoA of Dox (Fig. 3). Enriched GO BP terms included chromatin remodeling, regulation of transcription, DNA damage response (DDR), DNA repair, cell division, and translation, consistent with cellular responses to DNA intercalation and topoisomerase 2 inhibition [24]. Chromatin-remodeling terms were consistently downregulated in the nuclear fraction, whereas translation-related terms were consistently downregulated in the cytoplasmic fraction. The latter observation is in line with attenuation of translation during Dox-induced stress [27]. Overall, Dox treatment generated characteristic regulomes with shared enriched terms and similar directions of perturbation in cytoplasmic–nuclear response space. Although chromatin remodeling was enriched across all four Dox-treated cell lines, the underlying protein composition differed strongly. The number of regulated proteins assigned to this term ranged from 18 to 107 across cell lines (Fig. 3 B), with only limited overlap in contributing proteins (Fig. 3 C). Only CDK1 (P06493), a regulator of G2/M progression, and SMARCC1 (Q92922), a SWI/SNF chromatin-remodeling subunit, were shared across all four cell lines. This illustrates that shared enriched GO terms can arise from distinct protein-level responses, highlighting the value of linking each regulome feature to its contributing regulated proteins.

**Fig. 3:**
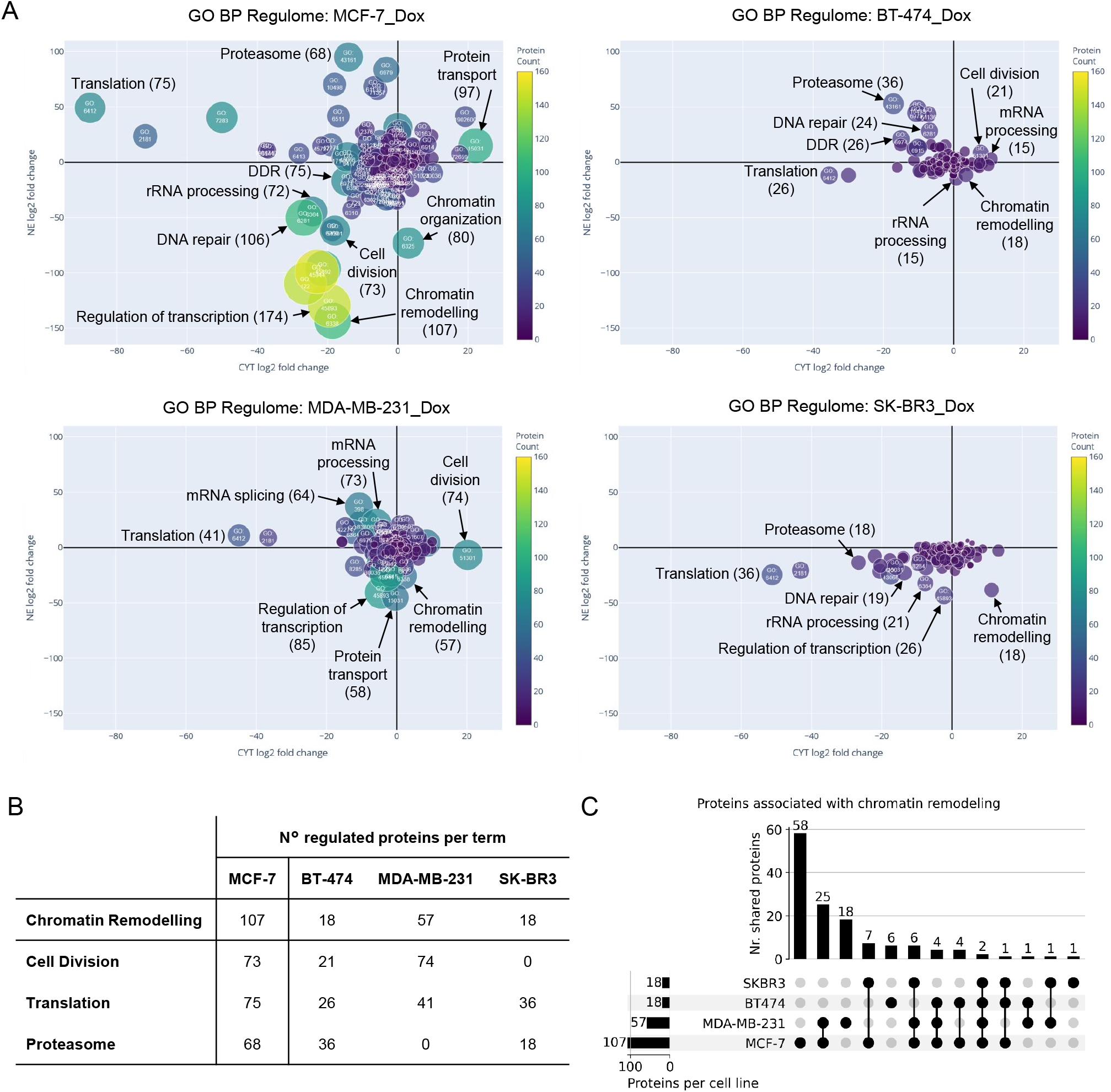
Comparison of the regulomes of Doxorubicin (Dox) treated breast cancer cells. (A) Regulomes of the four breast cancer cells treated with Dox, highlighting selected GO BP terms associated with the Dox response. (B) Table of the number of statistically significant protein regulations per term and cell line. (C) Upset plot showing the overlap of the regulated proteins in the term “chromatin remodeling” across the four breast cancer cell lines.

Among the four cell lines, MCF-7 (Luminal A) showed the strongest overall regulome perturbation after Dox treatment. This is notable because luminal breast cancer subtypes are generally considered less sensitive to Dox than HER2+ and triple-negative subtypes [23]. In addition, the two luminal cell lines, MCF-7 and BT-474, showed pronounced induction of proteasome-related terms in the nuclear compartment. Because proteasome-mediated degradation has been implicated in the removal of topoisomerase 2 cleavage complexes, and proteasome inhibition can potentiate topoisomerase 2-targeting drugs [28], this compartment-specific response may reflect a luminal-cell-line-associated stress adaptation to Dox treatment.

In contrast to the Dox response, Tax treatment produced a distinctly different regulome pattern centered on microtubule-associated processes, illustrated by the direct comparison of Dox- and Tax-treated MCF-7 cells (Fig. 4). In line with the microtubule-stabilizing MoA of Tax, the terms *microtubule cytoskeletal organization* and *microtubule-based process* were positioned in the nuclear-upregulated quadrant in three of the four cell lines (Supplementary Fig. S4). The strongest response was observed in MDA-MB-231 cells, followed by MCF-7 cells, whereas BT-474 showed a weaker but focused response dominated by these microtubule-related terms. By contrast, SK-BR3 showed only a weak overall response and did not display comparable microtubule-associated regulation.

**Fig. 4:**
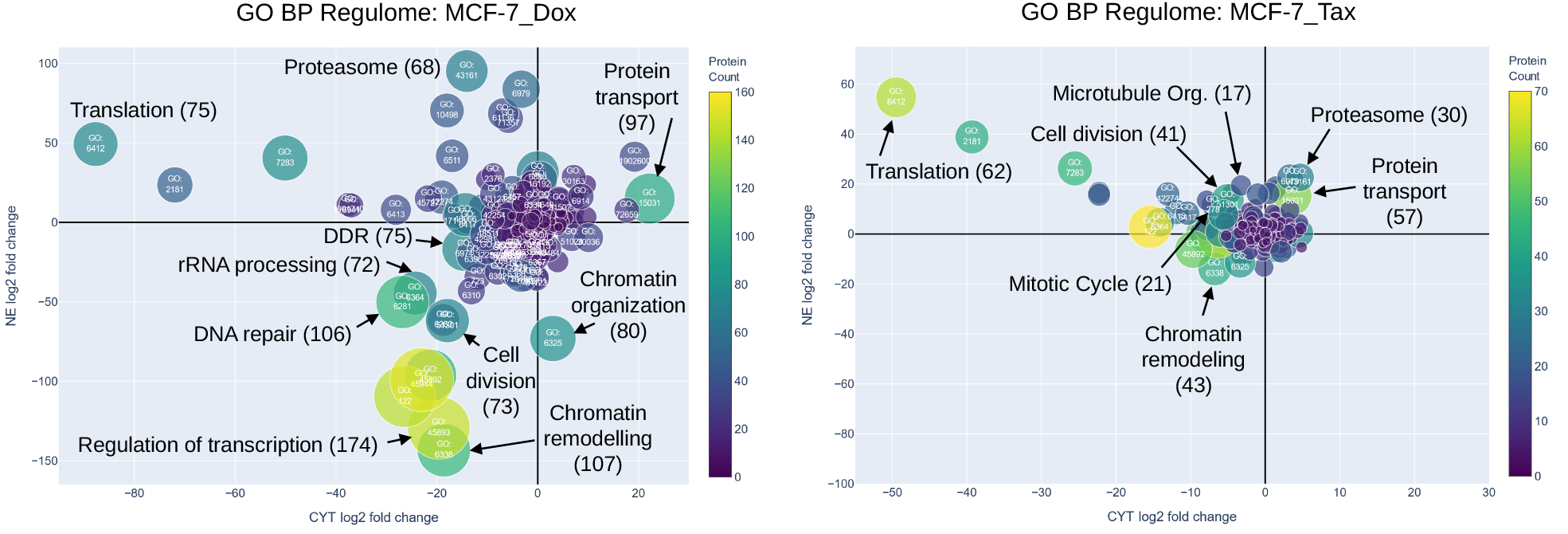
Comparison of drug-specific regulomes in MCF-7 breast cancer cells. Regulomes of MCF-7 cells treated with Dox (left) or Tax (right) show distinct compartment-resolved perturbation patterns. Selected GO BP terms associated with each drug response are highlighted. Note the different scales of axes and color bars of the two plots.

Similar to the chromatin-remodeling response after Dox treatment, the Tax-associated microtubule response was highly cell-line-specific at the protein level. The term *microtubule cytoskeletal organization* showed strong variation in the number of regulated proteins across cell lines and little overlap in contributing proteins (Supplementary Fig. S4 B and C, respectively). Thus, even when a microtubule-associated regulome feature was observed across multiple cell lines, it was driven by largely distinct sets of regulated proteins.

These examples demonstrate that regulomes enable an efficient comparison of perturbation profiles with subcellular resolution, revealing the number, magnitude and direction of potentially involved biological processes via GO term and pathway enrichment analysis, which in turn also facilitates discovering new aspects of drug MoA or resistance pathways.

## Code and data availability

RegulomeXplorer is freely available without registration at https://chemnettools.anc.univie.ac.at/RegulomeExplorer/. Its source code is available under the MIT license at https://github.com/chemnet-univie/regulome-xplorer.

Processed input data and example datasets used in this study are provided with the source-code repository.

Raw mass spectrometry data are available from the original studies cited in the manuscript.

## Author contributions statement

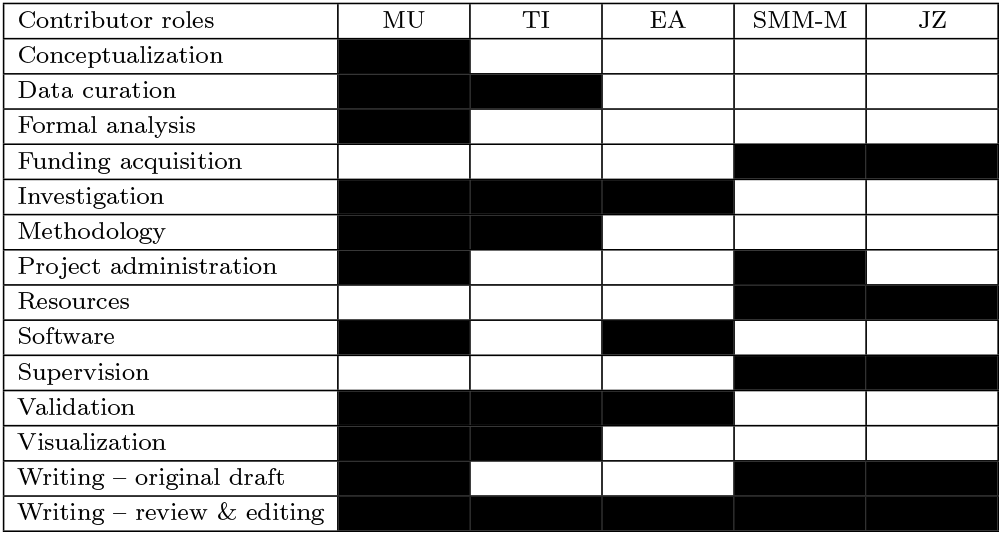

## Acknowledgments

The authors are grateful to the Core Facility for Mass Spectrometry and the Joint Metabolome Facility (University of Vienna and Medical University of Vienna), which are members of the Vienna Life-Science Instruments (VSLI). T.I. and S.M.M.-M. acknowledge financial s upport b y t he A ustrian S cience Fund (FWF), Grant doi:10.55776/PAT2536323.

