## Supplementary information for "RegulomeXplorer: Interactive exploration of drug effects on subcellularly resolved proteomes"

#### Input File Format

The workflow accepts protein group tables exported from MaxQuant in tab-separated text format (`proteinGroups.txt`). Each row corresponds to a protein group and columns contain protein annotations, identification metrics, peptide counts, sequence coverage statistics, and quantitative intensity measurements.

##### Required Columns

The current workflow requires the following columns for data import and analysis:

| Column | Description |
|---|---|
| `Protein IDs` | Uniprot protein accession numbers used as unique protein identifiers. |
| `Gene names` | Gene symbols associated with the identified protein. |
| `LFQ intensity <sample>` | Multiple columns with label-free quantification (LFQ) intensities for all samples. |

**Table S1.** Required columns for workflow execution.

The `Protein IDs` column is used as the primary identifier throughout the workflow. Here the first, most probable protein ID is used from the MaxQuant output. The corresponding `Gene names` are used for annotation, visualization and filtering. Sample replicates and conditions are automatically parsed from the `LFQ intensity <sample>` column headers. Differential expression analysis is currently performed exclusively on LFQ intensity values.

##### Quantitative Data

Quantitative measurements must be provided as individual columns following the MaxQuant naming convention:

```
LFQ intensity <sample_name>
```

For example:

```
LFQ intensity SW480_CYT_Con_1
LFQ intensity SW480_CYT_Con_2
LFQ intensity SW480_CYT_Re001_1
LFQ intensity SW480_NE_Re001_1
```

Each column represents one biological or technical replicate. Naming convention is `[column name, e.g. LFQ intensity]` followed by a space and then the following identifiers separated by underscores: cell-id, fraction (*CYT* or *NE*, which stands for cytoplasmic fraction or nuclear extraction, respectively), condition (*Con* for control or a treatment ID), biological replicate ID, and optional technical replicate ID.

##### Optional Metadata Columns

Additional columns present in the MaxQuant export are imported and stored in the database but are currently not required for statistical analysis. They might be used in future versions. These include, among others:

- `Majority protein IDs`
- `Protein names`
- `Fasta headers`
- peptide count metrics
- sequence coverage statistics
- molecular weight and sequence length information
- identification type annotations
- raw intensity values (`Intensity`)
- MS/MS counts
- peptide and evidence identifiers
- contaminant and reverse hit annotations

These metadata are retained to support future workflow extensions, additional quality-control procedures, and alternative quantification strategies.

##### Missing Values

Missing quantitative values should remain empty or contain the original MaxQuant missing-value representation. The workflow performs missing-value handling during downstream processing and therefore does not require prior imputation.

##### Filtering

The workflow automatically excludes entries marked as contaminants or reverse database hits when the corresponding MaxQuant annotation columns (`Potential contaminant`, `Reverse` and `Only identified by site`) are available. Entries are excluded if marked by a "+".

#### Regulome verification

To validate the automated workflow, we reanalyzed the manually curated nucleocytoplasmic proteomics dataset reported by Skos et al. [19] with our approach and compared the results with the published findings. The published workflow was performed in Perseus using left-censored imputation of the proteome of each fraction followed by differential expression analysis with the permutation-based SAM algorithm (Significance Analysis of Microarrays). As our workflow uses the limma package for differential expression by default which does not require imputation, we switched here to SAM from the *siggenes* package of Bioconductor, used the same parameters (FDR=0.05, $S_0$=0.1, Number of permutations=250) and loaded the original imputed data table as input into the workflow. The input table consisted of 6 samples per condition after imputation. Only proteins with a minimum of 5 measured values for at least one condition were accepted for imputation. The volcano plots obtained from both approaches were compared and show identical datapoints (see Supplementary Fig. S1). As both versions of SAM are not 100% identical, there is a slight difference in the position of the significance threshold curve which leads to a slightly different number of significant proteins with SAM of *siggenes* than originally published (see orange hyperbolic curve in Supplementary Fig. S1).

Representative abundance distributions for HMOX1 (P09601) and DNAJB1 (P25685) were examined to confirm consistent fold-change and significance estimates.

For both workflows, proteins passing multiple-testing correction were subsequently used as input for gene ontology enrichment analysis via the *David* web service. The resulting nucleocytoplasmic regulome plots for biological processes (GO BP) and cellular compartments (GO CC) generated by the

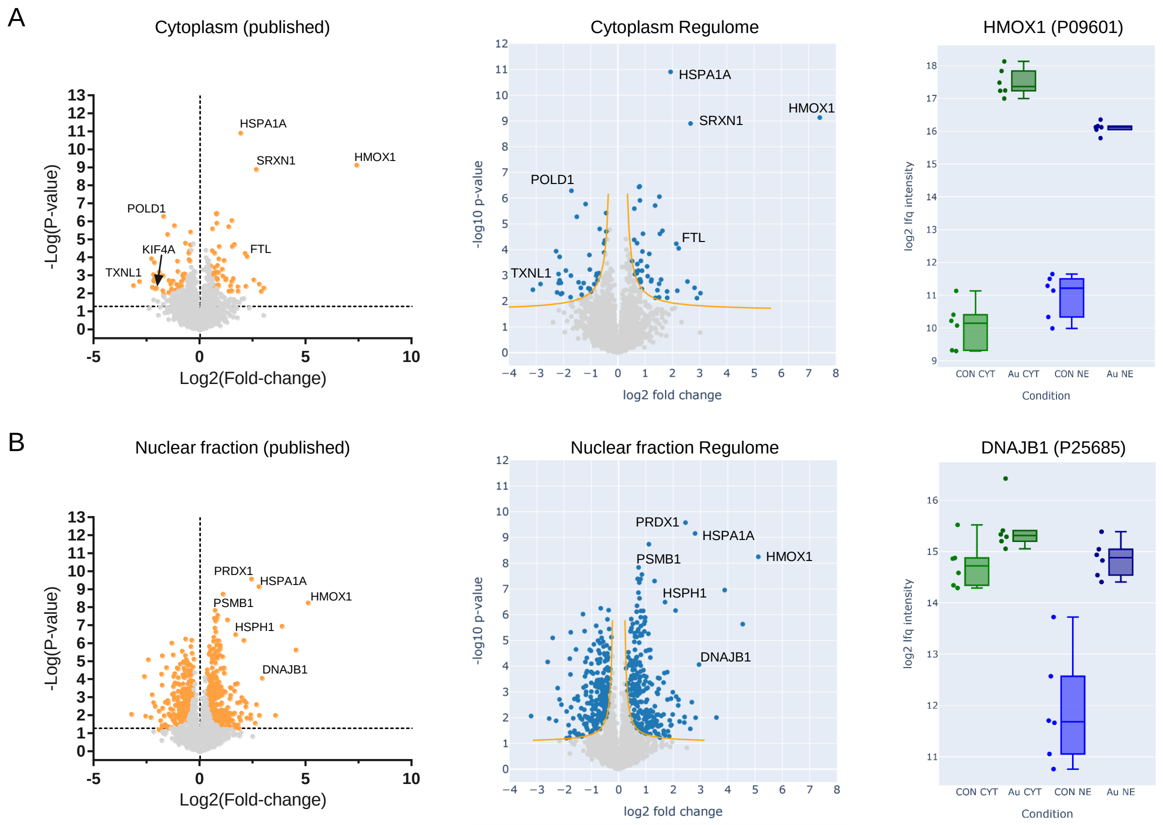


Supplementary Fig. S1: Verification of the automated data processing workflow against a manually curated regulome published in [19]. Comparison of volcano plots generated from published cytoplasmic (A) and nuclear (B) proteomics data using the original Perseus-based workflow and the automated workflow presented in this study. The orange curve represents the hyperbolic significance threshold of the SAM algorithm. Representative protein abundance distributions are shown for HMOX1 (A) and DNAJB1 (B).

automated workflow closely reproduced the manually curated reference results. The remaining differences stem from the differences in significant proteins and the different version of the *David knowledgebase* (currently v2026_1 from 23.5.2026) which is updated multiple times every year (see Supplementary Fig. S2).

### Regulome visualization mode

The regulome plot allows two different modes for data representation. In sum-mode the position of a term is determined by the sum of all fold-change values of the proteins within that term. Fold-changes of the cytoplasmic fraction are depicted on the x-axis and fold-changes of the nuclear fraction on the y-axis, respectively. This visualization has the advantage that larger terms are typically placed further away from the center which enables a good overview what the major relevantly regulated terms are (Supplementary Fig. S3 A).

In average-mode the position is determined by the average of all fold-change values of the proteins within that term (Supplementary Fig. S3 B). Here the terms are further from the center which have their proteins regulated unisono in the same direction, no matter how many proteins the term consists of. Here the small terms with less than 30 proteins were excluded for plotting to get a clearer picture for the larger terms.

For comparison, the term *translation* (GO: 6412) is highlighted and the corresponding protein distribution is shown in the inset of Supplementary Fig. S3 A. It mainly consists of proteins that are down-regulated in the cytoplasmic fraction and shown far left in both regulome plots. Notice that the term *cytoplasmic translation* (GO: 2181), which is in close vicinity, is right of it in sum-mode and left of it in average-mode. The reason here-fore is, that *cytoplasmic translation* is mainly a subset of *translation* but covers the more down-regulated proteins which push its average to the left in Supplementary Fig. S3 B. In contrast to this, *translation* is pushed more to the left in Supplementary Fig. S3 A due to its higher number of proteins.

Note also that some proteins of the protein distribution shown in the inset in Supplementary Fig. S3 A, are positioned exactly on either the CYT or NE axis. These are proteins where either the fold-change of the cytoplasmic or the nuclear fraction could not be calculated due to missing values.

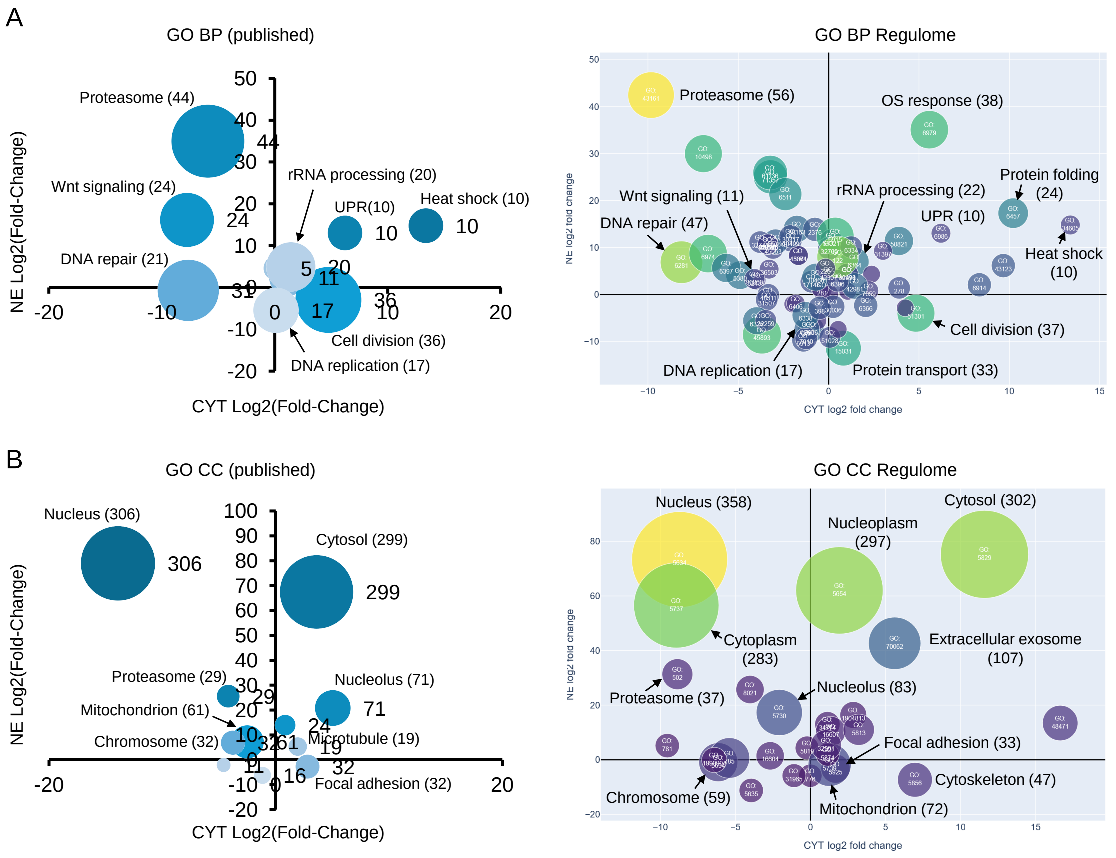


Supplementary Fig. S2: Comparison of manually curated regulomes from [19] and automatically generated nucleocytoplasmic regulome plots for gene ontology biological processes (GO BP; A) and cellular compartments (GO CC; B).

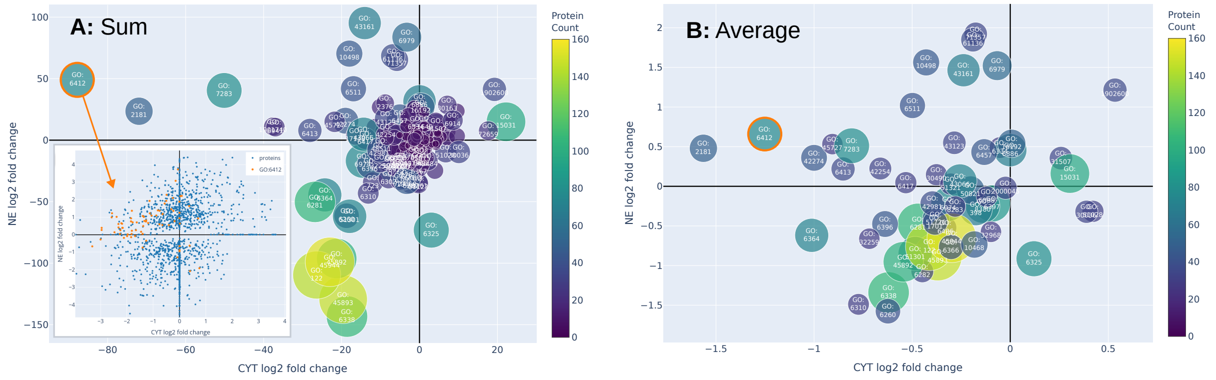


Supplementary Fig. S3: Comparison of the display modes of the regulome of biological processes for Doxorubicin (Dox) treated MCF-7 cells. (A) Sum-mode: For each GO term, the x (CYT) and y (NE) coordinates correspond to the cumulative fold-changes of all significantly regulated proteins assigned to that term in the CYT and NE fractions, respectively. (B) Average-mode: The average fold-changes for CYT and NE are calculated instead. Missing fold-changes due to missing LFQ intensities are ignored for sum and average calculations. The inset shows the 2D distribution of all significant proteins in blue and proteins of the term *translation* (GO: 6412) in orange.

A

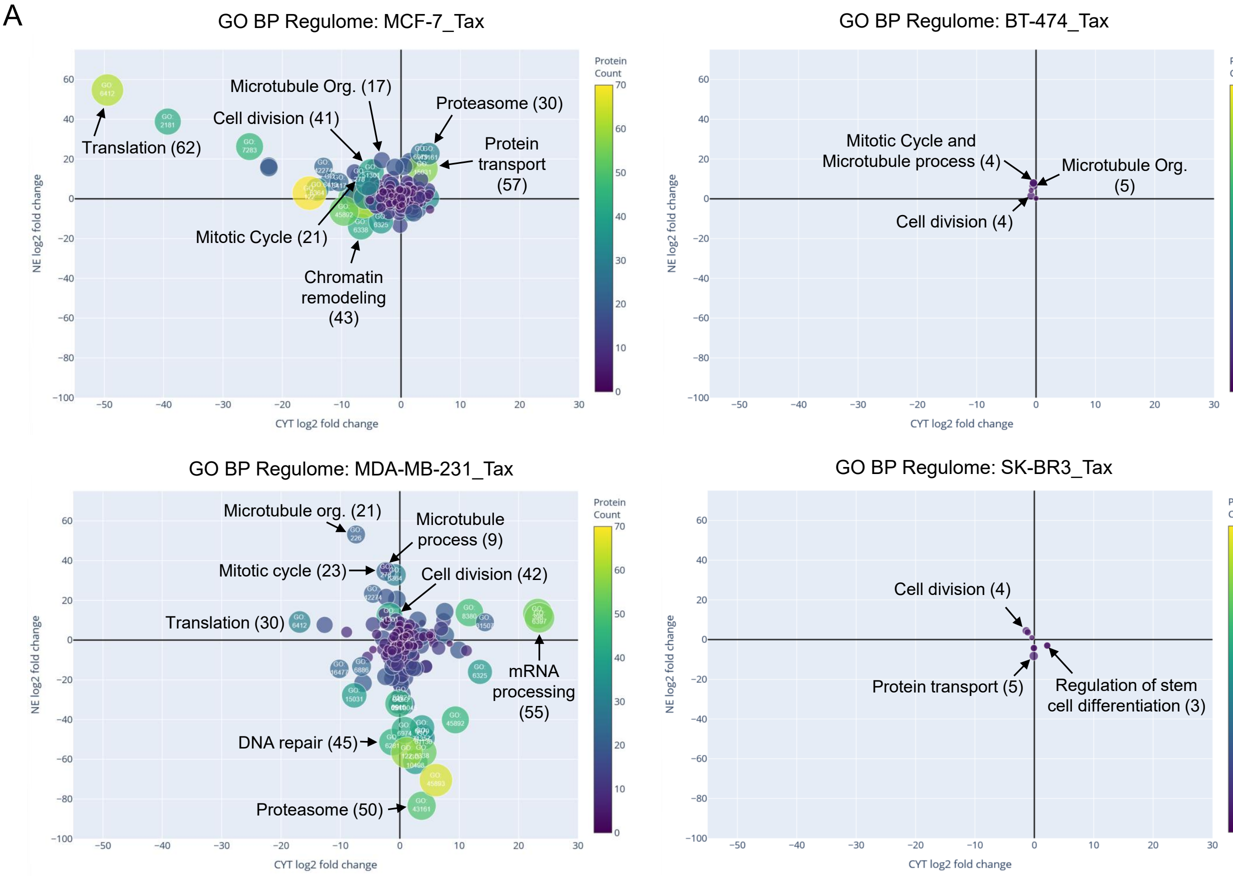


B

| | N° regulated proteins per term | | | |
|---|---|---|---|---|
| | **MCF-7** | **BT-474** | **MDA-MB-231** | **SK-BR3** |
| **Microtubule cytoskeleton organization** | 17 | 5 | 21 | 0 |
| **Microtubule process** | 9 | 4 | 9 | 0 |
| **Cell division** | 41 | 4 | 42 | 4 |
| **Mitotic cell cycle** | 21 | 4 | 23 | 0 |

C

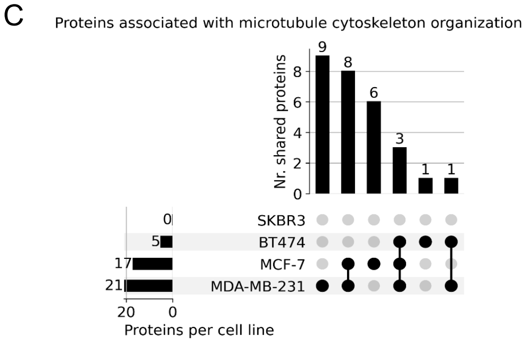


Supplementary Fig. S4: Comparison of the regulomes of Taxol (Tax) treated breast cancer cells. (A) Regulomes of the four breast cancer cells treated with Tax, highlighting microtubule-related and other relevant GO BP terms. (B) Table of the number of statistically significant protein regulations per term and cell line. (C) Upset plot showing the overlap of the regulated proteins in the term *microtubule cytoskeleton organization* across the four breast cancer cell lines.
